# Flame retardant tetrabromobisphenol A (TBBPA) disrupts histone acetylation during zebrafish maternal-to-zygotic transition

**DOI:** 10.1101/2024.03.31.587433

**Authors:** Rosemaria Serradimigni, Alfredo Rojas, Connor Leong, Uttam Pal, Madeline Bryan, Sunil Sharma, Subham Dasgupta

## Abstract

3,3’,5.5’-Tetrabromobisphenol A (TBBPA) is a widely used brominated flame-retardant utilized in the production of electronic devices and plastic paints. The objective of this study is to use zebrafish as a model and determine the effects of TBBPA exposure on early embryogenesis. We initiated TBBPA exposures (0, 10, 20 and 40μM) at 0.75 h post fertilization (hpf) and monitored early developmental events such as cleavage, blastula and epiboly that encompass maternal-to-zygotic transition (MZT) and zygotic genome activation (ZGA). Our data revealed that TBBPA exposures induced onset of developmental delays by 3 hpf (blastula). By 5.5 hpf (epiboly), TBBPA-exposed (10-20 μM) embryos showed concentration-dependent developmental lag by up to 3 stages or 100% mortality at 40 μM. Embryos exposed to sublethal TBBPA concentrations from 0.75-6 hpf and raised in clean water to 120 hpf showed altered larval photomotor response (LPR), suggesting a compromised developmental health. To examine the genetic basis of TBBPA-induced delays, we conducted mRNA-sequencing on embryos exposed to 0 or 40 μM TBBPA from 0.75 hpf to 2, 3.5 or 4.5 hpf. Read count data showed that while TBBPA exposures had no overall impacts on maternal or maternal-zygotic genes, collective read counts for zygotically activated genes were lower in TBBPA treatment at 4.5 hpf compared to time-matched controls, suggesting that TBBPA delays ZGA. Gene ontology assessments for both time- and stage-matched differentially expressed genes revealed TBBPA-induced inhibition of chromatin assembly- a process regulated by histone modifications. Since acetylation is the primary histone modification system operant during early ZGA, we immunostained embryos with an H3K27Ac antibody and demonstrated reduced acetylation in TBBPA-exposed embryos. Leveraging in silico molecular docking studies and in vitro assays, we also showed that TBBPA potentially binds to P300- a protein that catalyzes acetylation- and inhibits P300 activity. Finally, we co-exposed embryos to 20 μM TBBPA and 50 μM n-(4-chloro-3-trifluoromethyl-phenyl)-2-ethoxy-6-pentadecyl-benzamide (CTPB) -a histone acetyltransferase activator that promotes histone acetylation- and showed that TBBPA-CTPB co or pre-exposures significantly reversed TBBPA-only developmental delays, suggesting that TBBPA-induced phenotypes are indeed driven by repression of histone acetylation. Collectively, our work demonstrates that TBBPA disrupts ZGA and early developmental morphology, potentially by inhibiting histone acetylation. Future studies will focus on mechanisms of TBBPA-induced chromatin modifications.

## INTRODUCTION

Flame retardants (FRs) encompass various compounds designed to impede or suppress fire spread within furniture, electronics, vehicles, etc. Brominated flame retardants (BFRs) constitute a prominent class of FR chemicals that are of high production volume due to low cost and high efficacy. Of these, tetrabromobisphenol A (TBBPA) is the highest production-volume brominated flame retardant, widely used in the production of electronic circuit boards (Zhou et al., 2014). TBBPA is a high priority chemical undergoing risk evaluations under Toxic Substances Control Act (TSCA) and identified as a potential carcinogen in California Proposition 65. While TBBPA exhibits remarkable FR properties, it is an additive FR and has the potential to leach out of its end use products, leading to its detection in various environmental and biological matrices, including indoor environments (Colnot et al., 2014). Although TBBPA and its conjugates have a half-life of 2-3 days in humans (Alexander et al., 2011; Cariou et al., 2008), chronic exposure to the parent chemical will continue to prevail for years from older end-use products within indoor environments. Within indoor environments, exposure to TBBPA can occur through inhalation, ingestion, or dermal contact (Lee et al., 2020; Oral et al., 2021). Over the years, significant parent and metabolite levels TBBPA has been detected in urine samples, placental tissue, breast milk and cord blood, confirming ubiquitous in utero exposures posing health risks. Indeed TBBPA has been detected adult blood and urine samples, as well as cord blood (Cariou et al., 2008), breast milk (Shi et al., 2009), and placental tissue (Alzualde et al., 2018). In fact, TBBPA concentrations upto 20 nM in adult plasma (Alzualde et al., 2018) and 5.3 nM in cord plasma (Sunday et al., 2022) have been detected, with a study modelling that concentrations up to 1.2 µM may be present in cord plasma (Alzualde et al., 2018). Detection of TBBPA in the latter group of biomatrices establish its potential to induce health effects in a developing embryo (Reed et al., 2022). Therefore, comprehensive evaluation of chemico-biological interactions is therefore necessary to minimize risk, formulate preventative and therapeutic strategies from current and future exposures and strategize remedial measures.

Over the past decade, numerous studies have identified TBBPA as a developmental toxicant, with targeted effects linked to reproductive and developmental adverse effects (Zhou et al., 2014), neurotoxic outcomes (Cho et al., 2020), and endocrine disruption (Oral et al., 2021). However, a critical window of development that has not been well-explored is the pre-pluripotent window, during with the zygotic genome is established. During these stages, the embryo undergoes a maternal-to-zygotic transition, during with the maternal transcripts degrade and the zygotic genome is (ZGA), shifting the embryo’s dependency on maternal RNA and proteins toward activating its own genetic program (Tadros & Lipshitz, 2009). This critical function, conserved across most species, is essential for controlling the advancement of developmental regulation. These developmental events are regulated by various genetic and epigenetic modifiers, including histone modifications that remodel the chromatin for transcription (Schulz & Harrison, 2019). The complexity of these processes makes them susceptible to xenobiotic exposures during these stages and any genetic and epigenetic interference can disrupt ZGA, impacting the downstream embryogenesis trajectory. Indeed, prior work has shown that drugs such as niclosimide can disrupt ZGA and delay zebrafish gastrulation (Vliet et al., 2018), indicating that external chemical stressors can potentially target these sensitive early embryogenesis stages.

Within this study, we used zebrafish as a model to evaluate impacts of TBBPA exposures on ZGA. Zebrafish was chosen as a model organism due to their genetic similarity to humans and their suitability for studying embryonic development, facilitating an evaluation of the effects of TBBPA exposure on ZGA. Specifically, the pre-pluripotent stages can be microscopically visualized *in vivo* for phenotypic effects as well as effects on specific gene products. Initially, phenotypic profiling revealed a significant delay in embryonic development induced by TBBPA exposure, particularly during the critical periods of maternal-to-zygotic transition (MZT) and zygotic genome activation (ZGA). We then investigated mechanisms of developmental delays by relying on-1) mRNA-sequencing to interrogate progression of ZGA and altered genetic pathways, followed by 2) a combination of pharmacological strategies, whole-mount immunohistochemistry, *in vitro* assays and molecular docking studies to examine how TBBPA targets a specific histone modification system-histone acetylation.

## METHODS

### Animals

Founder adult wild-type (strain 5D) zebrafish were obtained from Sinnhuber Aquatic Research Laboratory in Corvallis, OR and maintained on a recirculating system and housed within the Aquatic Animal Research Laboratory at Clemson University. Fish were kept under a 14:10 -h light-dark cycle at a water temperature of ∼28.5 ºC, pH of approximately 7-8, and conductivity levels of roughly 200-700 μS. Water quality was monitored manually for temperature and conductivity daily. Ammonia, nitrate, nitrite, alkalinity, and hardness levels were manually monitored using colorimetric strips. Zebrafish were fed twice daily with specialized adult zebrafish food (Zeigler Adult 1mm pellets, Zeigler Bros, Inc. 400 (Gardners Station Road, Gardners, PA 17324 USA). Zebrafish were bred off-system using static-renewal water within a breeding container with a false bottom insert. One adult male and female fish were kept overnight in the spawning chamber. For all experiments described, newly fertilized eggs were collected within 30 minutes of spawning, rinsed, and reared in a temperature-controlled incubator (28ºC). All embryos were sorted and staged according to previously described methods (Kimmel et al., 1995). All adult breeders were handled and treated per an Institutional Animal Care and Use Committee-approved animal use protocol (No. 20220434) at Clemson University.

### Chemicals

TBBPA (97% purity) (CAS #: 79-94-7) was purchased from Sigma-Aldrich. N-[4-chloro-3-(trifluoromethyl)phenyl]-2-ethoxy-6-pentadecyl-benzamide (CTPB) (CAS #: 586976-24-1) (>99% purity) was purchased from Abcam (Cambridge, United Kingdom). For both chemicals, stock solution was prepared in high-performance liquid chromatography-grade dimethyl sulfoxide (DMSO) and stored within 20 ml glass vials with parafilm-wrapped lids at room temperature. Working solutions were prepared in system water prior to each experiment by preparing a stock working concentration and serial dilution of the stock concentration into subsequent concentrations.

### Embryo Exposures and Phenotyping

All embryos were staged according to previously described methods (Kimmel et al., 1995). Viable embryos at the 2-cell stage (0.75 hours post fertilization (hpf)) were sorted into 4 replicates of 10 embryos within 24-well plates (VWR international, LLC Radnor, PA, USA). Replicates were exposed to 1 mL of vehicle (0.1% DMSO; henceforth referred to as 0 μM TBBPA) or various TBBPA concentrations. Experiments were conducted under static conditions at 28ºC within a temperature-controlled incubator for the duration of the exposure. Phenotypes were observed and imaged under an Olympus TH4-100 equipped with a Lumenera Infinity 8-8 camera (Ottawa, Ontario, Canada). Phenotyping of embryos included assessments of mortality, deformity and developmental stages for each embryo based on Kimmel et al 1995.

### Window of sensitivity experiments

To identify a potentially critical period where TBBPA impacts zebrafish embryogenesis, embryos (4 replicates of 10 embryos in 24-well plates) were treated with 0 or 40 μM TBBPA. Exposures were staggered in 1.15-hour intervals beginning at 0.75 hpf and all exposures terminating at 6 hpf. At each exposure time point (0.75, 2, 3.15, and 4 hpf), embryo media was removed from corresponding wells and 1 mL of TBBPA exposure solution was added to each well. Following termination of the experiment, embryos were phenotyped as described in previously.

### Embryo Recovery and Larval Photomotor Response (LPR)

To compare the developmental impacts of short -term TBBPA exposures, embryos (32 replicate embryos in 96-well plates) were exposed to 0, 0.004-, 0.04-, 0.4-, and 4 μM TBBPA (16 embryos per treatment group per concentration in a plate, 2 plates per experiment). We did not include the 40 μM concentration since this leads to mortality by 6 hpf. Exposures lasted from 0.75-6 hpf, following which the TBBPA solution was removed from each well, washed 3X and embryos were subsequently placed in chemical-free water from 6-120 hpf. Plates were kept in a dark box within a temperature-controlled incubator at 28°C. At 120 hpf, larval photomotor response of embryos was assessed on a Zebrabox (Viewpoint, France) over 4 cycles of 3 min lights on and 3 min lights off per cycle (epoch), and a 2500 lux intensity of light. Data was acquired using Viewpoint’s Zebralab software and analyzed in R (https://www.r-project.org/) using custom codes modified from previous work (Dasgupta et al., 2022). Dead and deformed embryos were excluded from analysis. Raw movement data for LPR is presented within Supplemental Table S1 and S2. Data interpretation was conducted based on 3^rd^ light (time bin 121-150) and dark (time bin 151-180) cycles from each run.

### mRNA Sequencing

To evaluate impacts on the transcriptome, embryos (20 per well in 24-well plates) were exposed to vehicle 0 or 40 μM TBBPA from 0.75 – 2 or 3.5 or 4.5 hpf and incubated at 28ºC. At each time point, 50 embryos per replicate were collected into RINO 1.5 mL screwcap tubes, homogenized in RNAzol using a Bullet Blender Storm Pro (Next Advance, Inc., Troy, NY, USA) and stored at -80ºC resulting in a total of 24 samples. Following homogenization, Direct-zol RNA MiniPrep Kit (Zymo Research Corporation, Irvine, CA, USA) was used to extract total RNA using the manufacturer’s instruction. mRNA quality and quantity were confirmed using a Agilent Tape station 4200 (Agilent, Santa Clara, CA) and Nanodrop (Thermo Scientific, Waltham, MA) and sent to Novogene Corporation Inc. for library preparation and mRNA-seq (see Supplemental File for RNA-seq details). Sequencing was conducted on a Novaseq 600 (Illumina, San Diego, CA) (2x150 bp, ∼40m reads/sample). Raw sequencing files will be deposited into NCBI Gene Expression Omnibus under accession # XXXX.

### Bioinformatic analysis of sequencing data

Raw data (raw reads) in fastq format was firstly processed through Novogene’s in-house perl scripts. In this step, clean data (clean reads) was obtained by removing reads containing adapter, reads containing poly-N and low-quality reads from raw data. Reads were mapped to reference genome (GRCz 11) using Hisat2 v2.0.5 . FeatureCountsv1.5.0-p3 was used to count the reads numbers mapped to each gene. Then FPKM (Fragments Per Kilobase of transcript sequence per Millions base pairs sequenced) of each gene was calculated based on the length of the gene and reads count mapped to this gene. Differential expression analysis of two conditions/groups was performed using the DESeq2R package (1.20.0). The resulting P-values were adjusted using the Benjamini and Hochberg’s approach for controlling the false discovery rate. Genes with an adjusted P-value <0.05 found by DESeq2 were assigned as differentially expressed. Following this, gene ontology assessment (biological processes) was conducted within ShinyGO v0.77. Expanded details for RNA-seq are provided within Supplemental File. Raw data for counts, differentially expressed genes and gene ontology are included within Supplemental Tables S3-S8.

### Assessments of maternal-to-zygotic transition

We parsed out normalized FPKM read counts for maternal (M), maternal-zygotic (MZ) and zygotic (Z) genes across the 3 sequencing time points to represent gene expression of these transcripts. transcripts. Information of these transcripts was obtained from a previous work (Bhat et al., 2023) and are listed in Supplemental Table S9. We plotted (FPKM + 1) values for each gene and grouped them into M, MZ and Z; 1 was added to FPKM values to represent data on a logarithmic scale.

### Whole-mount immunohistochemistry (IHC)

To examine and quantify TBBPA’s potential effect on histone acetylation, embryos (20 per concentration) were exposed to vehicle (0.1% DMSO), 5, 10, 20, or 40 μM TBBPA from 0.75-3.5 hpf. Embryos were then fixed in 4% paraformaldehyde overnight and immunohistochemistry was performed according to previous protocols (Dasgupta et al., 2019) with minor modifications using an anti-acetyl H3K27 – primary antibody (Abcam, AB177178; 2 µg/mL) and a goat-anti-rabbit IgG secondary antibody (Sigma Aldrich, 1:500 dilution). A separate pool of embryos was stained only with secondary antibody for measuring background fluorescence. Following immunostaining, embryos were imaged using ECHO Revolve R4 (San Diego, CA, USA) with Olympus 2X objective lens. Fluorescence was quantified using Image J as mean gray intensity of the cell mass for each embryo and adjusted for background fluorescence by subtracting the mean fluorescence of secondary antibody-only stained embryos.

### Molecular docking studies

To examine if TBBPA interacts with P300- a protein that catalyzes the transfer of acetyl groups from acetyl CoA onto chromatins, we first conducted molecular docking studies and assessed binding of TBBPA onto the catalytic domain and bromodomain of P300. All the available crystal structures information of p300 (UniProtKB: Q09472) comprising catalytic and/or bromodomain were obtained from the Protein Data Bank in Europe and/or PBD-REDO repositories (accessed 2023-06-04) and listed in Table S10. Model geometry, fit model/data, PDB-REDO model geometry and PDB-REDO fit model/data of the crystal structures were scored on an arbitrary scale of -2 (worst) to +2 (best) (Chakraborty et al., 2020) based on the wwPDB validation report (Smart et al., 2018). Crystallographic information of the best quality structures were obtained for molecular docking simulation. Chemical structure of the small molecules were obtained from PubChem: Tetrabromobisphenol A (Compound ID: 6618), Cholera Toxin B Subunit (Compound ID: 729859), GNE-781 (Compound ID: 132275066), CTPB (Compound ID: 10311918). Prior to docking, all the obtained protein crystal structures were aligned and bound ligands were removed. Crystallographic waters and other heteroatoms were also removed. Less probable alternate positions of the side chains were removed. Missing residues were modelled in PyMOL. Molecular docking was performed using AutoDock Vina (Trott & Olson, 2010). Protein and ligand structures were prepared for docking using AutoDockTools (Morris et al., 2009). During structure preparation for docking, polar hydrogens were added to the protein structures and non-polar hydrogens were removed from the ligand structures; rotations were allowed on all the rotatable bonds of the ligands. Binding sites were defined based on the known bound ligands (bound ligand information is listed in Supplemental table S10). Docking simulations were run targeting the catalytic domain as well as the bromodomain. Lowest possible binding energies obtained from docking are listed in Supplemental Table S11-S12.

### In vitro P300 Assay

To quantify the potential for TBBPA to inhibit P300 *in vitro*, we used a SensoLyte® HAT (p300) Assay Kit (Anaspec, Fremont, CA). Assay was conducted on a black walled 96-well petriplate (VWR) according to manufacturer’s instructions and fluorescence was measured on an Agilent BioTek Microplate Reader (Agilent, Santa Clara, CA) at 390 excitation /513 emission wavelengths.

### CTPB co-exposure & pre-exposure

To assess the role of P300 in driving TBBPA-induced phenotypes, we conducted both pre-exposure and co-exposure experiments with TBBPA and CTPB- a P300 activator. For co-exposures, embryos were co-exposed to 0 and 20 µM TBBPA in presence or absence of 50 µM CTPB (4 replicates of 10 embryos in 24-well plates) from 0.75-3 hpf. For pre-exposure experiments, embryos were exposed to 0 or 50 µM CTPB from 0.75-2 hpf, followed by 0 or 20 µM TBBPA from 2-5.5 hpf (4 replicates of 10 embryos in 24-well plates). The change in phenotyping time point between co- and pre-exposures was done to provide sufficient time for both CTPB and TBBPA exposures. Only 20 µM TBBPA was chosen to avoid TBBPA-induced mortality at 5.5 hpf. Following termination of the experiment, embryos were phenotyped as described in “Embryo Exposures and Phenotyping”.

### Statistics

All statistical analyses and generation of images were performed within GraphPad Prism 9.0.

#### Initial phenotyping

A 2-way ANOVA (concentration x stage), followed by Dunnett’s posthoc test (p<0.05) was used to compare overall concentration x stage interaction and pairwise comparisons of each concentration to controls at each individual stage.

#### Window of sensitivity

A 2-way ANOVA (window x stage), followed by Tukey’s posthoc test (p<0.05) was used to compare overall window x stage interaction and pairwise comparisons of all windows to each other at each embryonic stage.

#### LPR assay

Data from epoch 3 were used for statistical estimations. Statistics for light and dark periods were calculated separately. Statistical differences were estimated based on a one-way repeated measures ANOVA, followed by a Dunnett’s test (p<0.05).

#### Maternal-to-zygotic transition

Since the 2 hpf timepoint was a separate sequencing experiment, a Kruskal-Wallis test (p<0.05) was conducted to identify statistical significance of the M, MZ or Z read counts at that time point. For the 3.5 and 4.5 hpf timepoints, a 1-way ANOVA was used, followed by a Tukey’s posthoc test (p<0.05).

#### Immunohistochemistry

A 1-way ANOVA, followed by a Dunnett’s posthoc test (p<0.05) was used to compare overall concentration effects and pairwise comparisons of each concentration to controls.

#### In vitro P300 assay

A generalized linear regression model was used to assess the overall effects of TBBPA exposure (p<0.05).

#### Co- and pre-exposure experiments

A 3-way ANOVA was used with stage, TBBPA and CTPB as factors, followed by Tukey’s posthoc test. For the stage factor, we chose to calculate the number of embryos at a specific stage, rather than assessments of all stages. For co-exposure experiments, stage includes presence or absence of embryos at the 128-256 cell stage. For pre-exposure experiments, stage includes presence or absence of embryos at the sphere cell stage.

## RESULTS

### TBBPA induces concentration-dependent developmental delays during ZGA window

We initiated embryonic exposures to TBBPA (0-40 µM) at 4-cell stage and phenotyped at various time points till 5.5 hpf **(Fig 1A)**. At ∼3 hpf, control embryos appeared in the high-oblong stage of development, 100% treated embryos displayed ∼0.75-1 hour delay in development. Furthermore, embryos displayed ∼2 hour concentration-dependent delay or mortality by 5.5 hpf. Statistical analyses showed that at every time point, stage x concentration interactions were statistically significant. There was a concentration-dependent difference (delay) in number of embryos compared to the control at a specific time point. For eg, at 5 hpf, while control embryos were at ∼50% epiboly stage, TBBPA-treated embryos ranged from high to dome stages depending on their concentrations. Our window of sensitivity experiment revealed an overall window x stage effect (p<0.0001) and that the sensitive window for mortality effects lay between 2 and 3.15 hpf, overlapping with the onset of ZGA **(Fig 1B)**; while exposures initiated at 0.75 and 2 hpf showed 0% survival at 5.5 hpf, subsequent exposures initiated at 3.15 and 4.5 hpf displayed ∼90% and ∼95% survival at 5.5 hpf, respectively.

**Figure 1.**
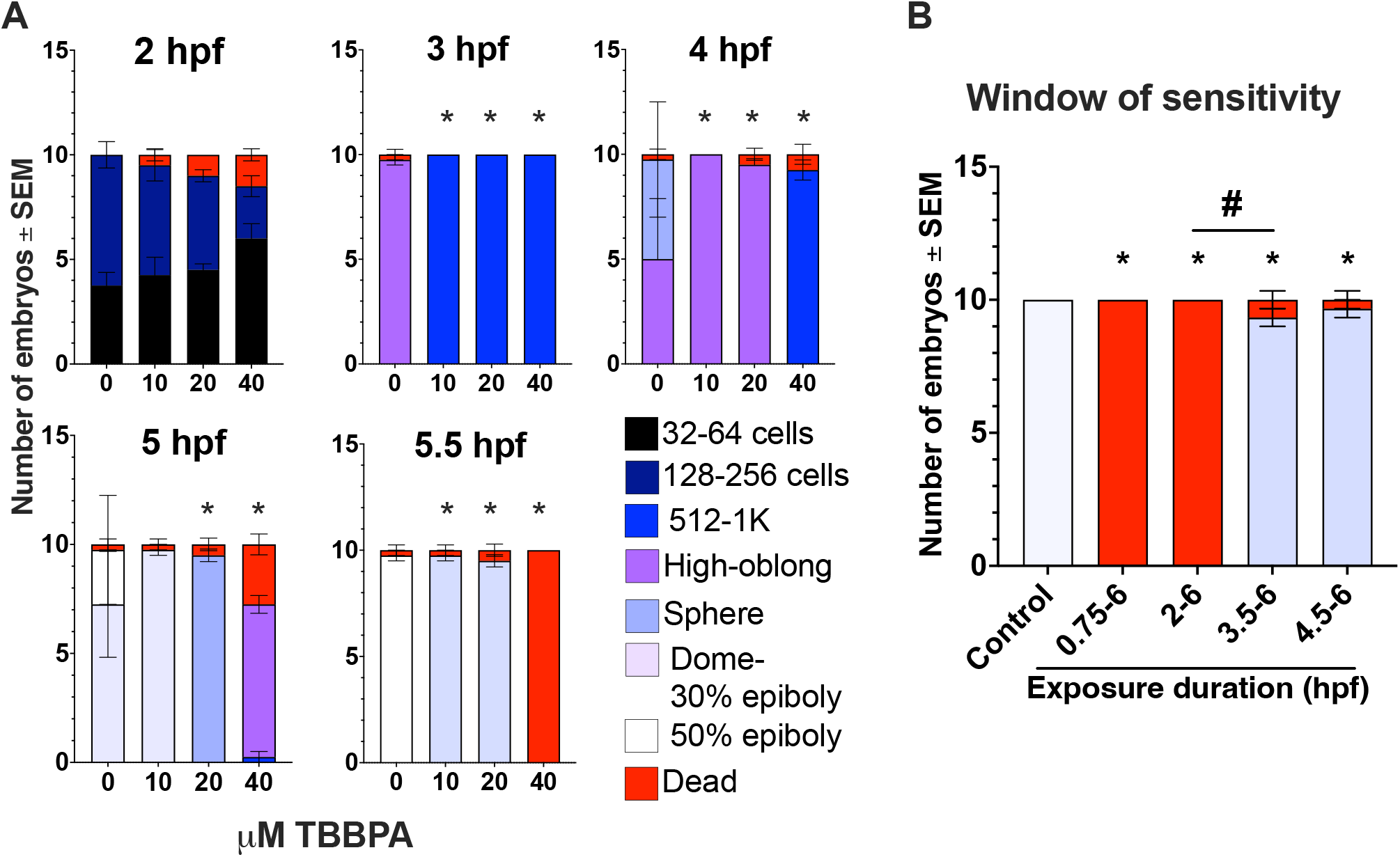
TBBPA exposures cause developmental delays. **(A)** Initiation of TBBPA exposure at 0.75 hpf induces concentration-dependent developmental delay starting at 3 hpf. **(B)** Exposures beginning prior to 3.15 hpf result in 100% mortality, whereas exposures following this time show significant survival. Asterisk (*) denotes statistically different from 0 µM based on 1-way ANOVA followed by Dunnett’s test. Pound sign (#) denotes 2-6 exposure duration significantly different from 3.5-6 hpf.

### TBBPA exposures induce hypoactivity during LPR at environmentally relevant concentrations

Our LPR data showed significant decrease (p<0.05) in LPR following exposure to TBBPA at environmentally relevant concentrations (< 0.1 µM); larvae exposed to TBBPA exhibited hypoactivity across both light and dark cycles **(Fig 2)**. This suggests that short-term exposures to TBBPA can have long-term effects on developmental health.

**Figure 2.**
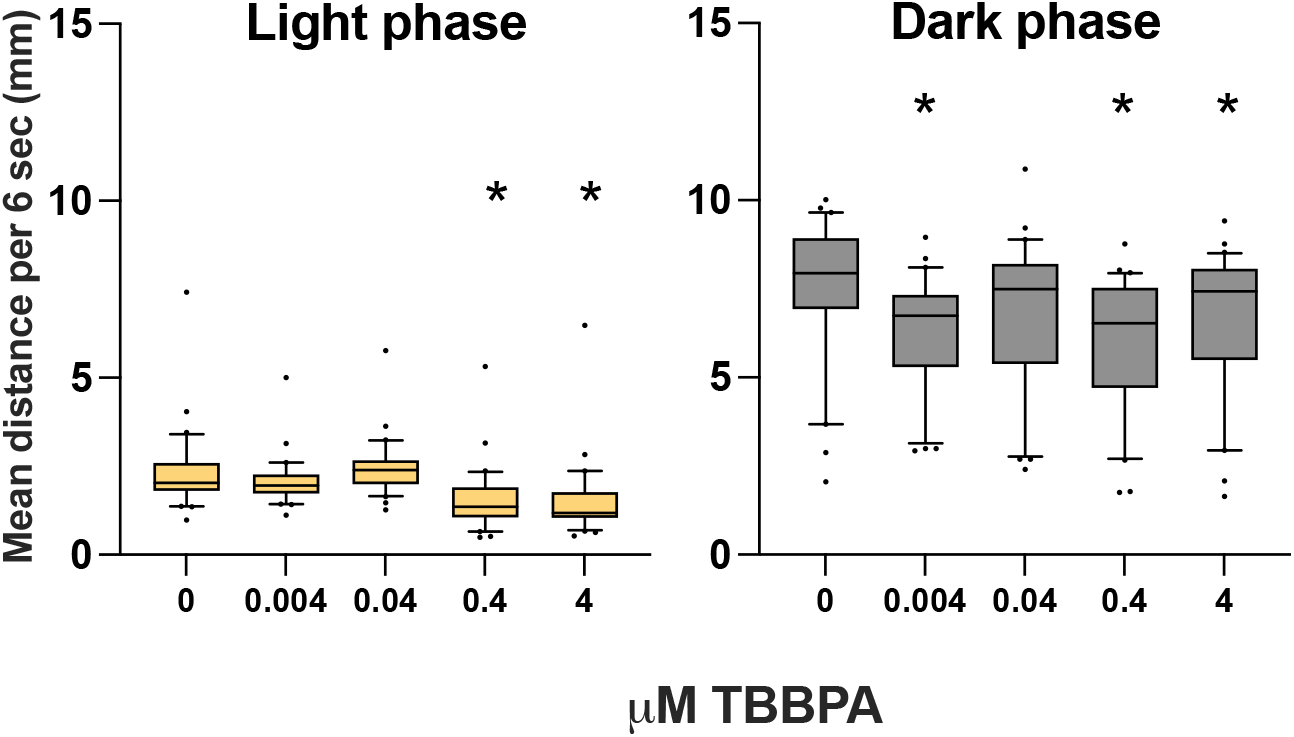
TBBPA exposures induce hypoactivity during larval photomotor response. TBBPA exposure from 0.75 to 6 hpf, followed by recovery in chemical-free water from 6 – 120 hpf cause hypoactivity in both light and dark phases. N=32 embryos in 96-well plates. *statistically different from 0 µM based on repeated measures 1-way ANOVA followed by Dunnett’s post hoc test (p<0.05) for 3rd light and dark cycles.

### mRNA sequencing shows that TBBPA disrupts ZGA and chromatin remodeling

To determine genetic targets of TBBPA, we conducted mRNA sequencing on embryos exposed to 0 or 40 µM TBBPA from 0.75-2, 3.5 or 4.5 hpf (40M reads/sample, N=4). We plotted FPKM values for maternal (M), maternal-zygotic (MZ) and zygotic (Z) transcripts; list of these transcripts was obtained from previous work (Zhang et al., 2018). While collective read counts of M and MZ transcripts were not affected, zygotic transcripts for TBBPA 4.5 hpf had significantly lower read counts than DMSO-4.5 hpf with a ∼2-fold decrease, but comparable to DMSO 3.5 hpf **(Fig 3)**. For differential gene expression, we performed *time-matched* and *stage-matched* comparisons and applied a |log_2_FC|>1 and p_adj_ <0.05 as cutoffs. Time-matched comparisons between DMSO and TBBPA at 4.5 present 3,294 differentially expressed genes while stage-matched comparisons of 3.5 hpf DMSO embryos against 4.5 TBBPA embryos revealed 261 differentially expressed genes **(Fig 4A)**. Gene Ontology revealed several pathways that were impacted **(Table S5-S8)** but the top inhibited pathways (based on GO of negative log fold change values; **Table S6, S8**) were associated with chromatin remodeling and nucleosome assembly **(Fig 4B)**.

**Figure 3.**
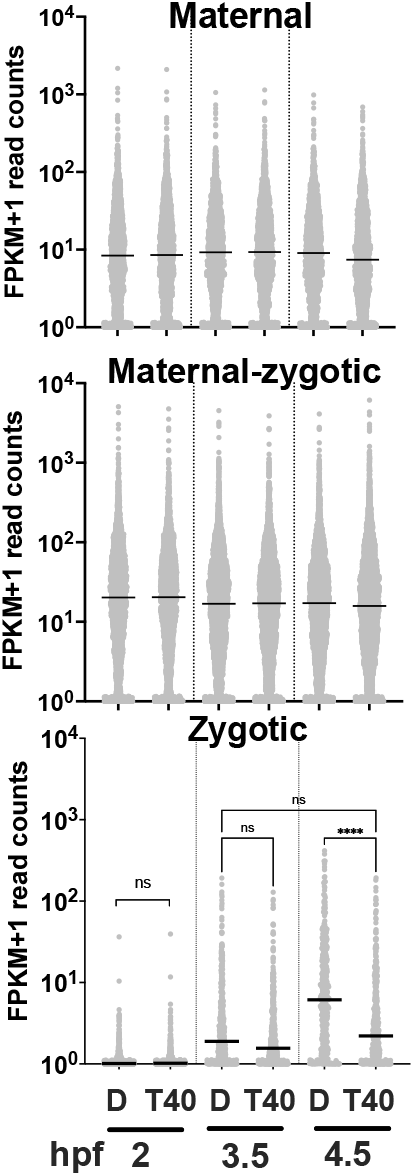
mRNA sequencing shows that TBBPA reduces mRNA levels of zygotic transcripts but not maternal-only or maternal-zygotic transcripts. Exposures from 0.75 to 2, 3.5 or 4.5 hpf. Asterisk (*) denotes statistically different based on 1-way ANOVA followed by Tukey’s test (p<0.05). D= DMSO, T40= 40 µM TBBPA.

**Figure 4.**
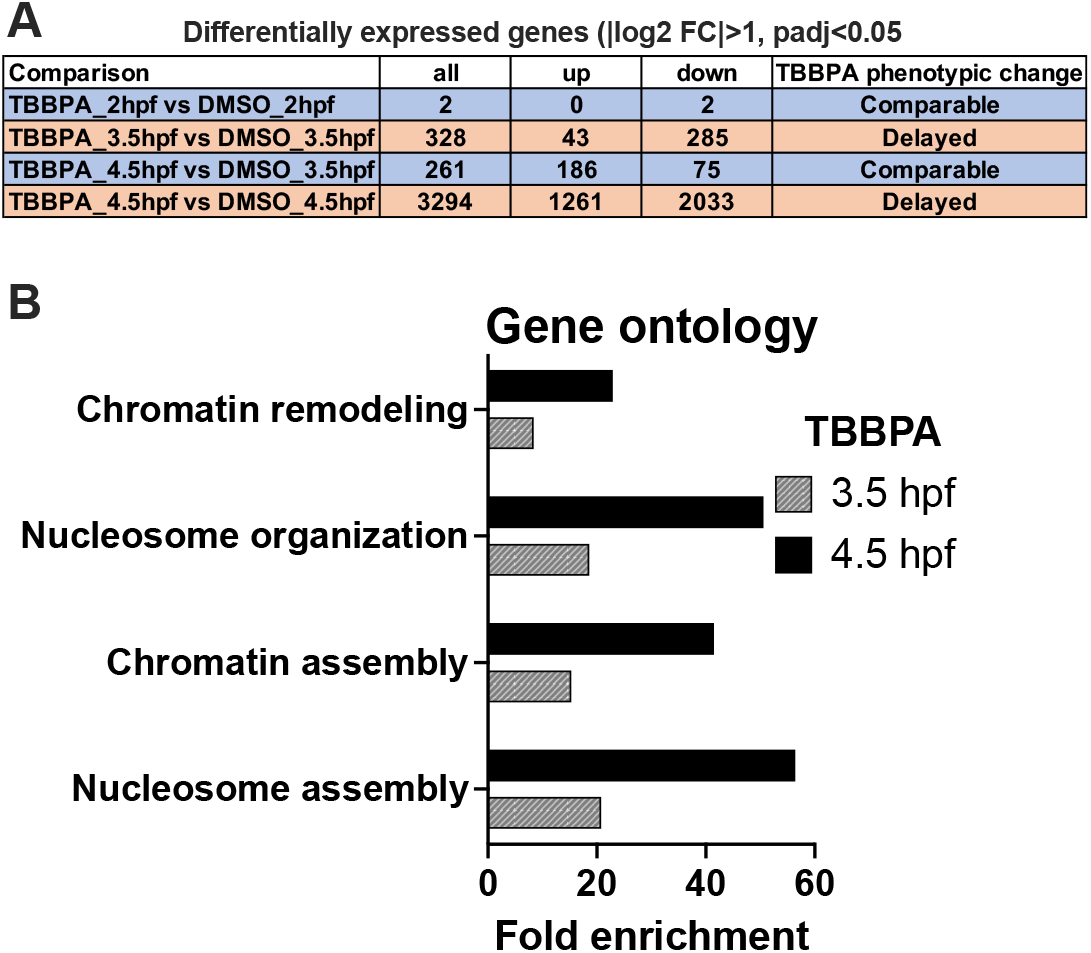
TBBPA disrupts expression of genes associated with chromatin remodeling. **(A)** Number of differentially expressed genes in time- or stage-matched comparisons. **(B)** Gene ontology (Biological Processes) reveals that TBBPA inhibits various pathways regulating chromatin assembly both for time and stage matched embryos. See full dataset in Tables S6 and S8.

### TBBPA inhibits H3K27Ac levels in vivo and P300 protein

Based on our sequencing data, we sought to evaluate of TBBPA inhibits H3K27 acetylation (H3K27Ac)- a histone modification system that remodels chromatin and is dominantly operant during embryonic stages where we see TBBPA-induced phenotypes. Using IHC, we quantified the extent of H3K27 acetylation within developing embryos. Overall, H3K27Ac levels showed a significant concentration-dependent reduction within the cell mass, with a ∼50% reduction in 20 µM TBBPA treatments and ∼75% reduction in 40 µM TBBPA compared to controls **(Fig 5)**. Our *in silico* molecular docking assessments showed that binding energies of CTPB and TBBPA to P300 catalytic domain are comparable **(Fig 6A)**. Finally, in concordance with our *in vivo* data, our *in vitro* P300 assay showed a P300-induced inhibition of P300 activity (p=0.0024) **(Fig 6B)**.

**Figure 5.**
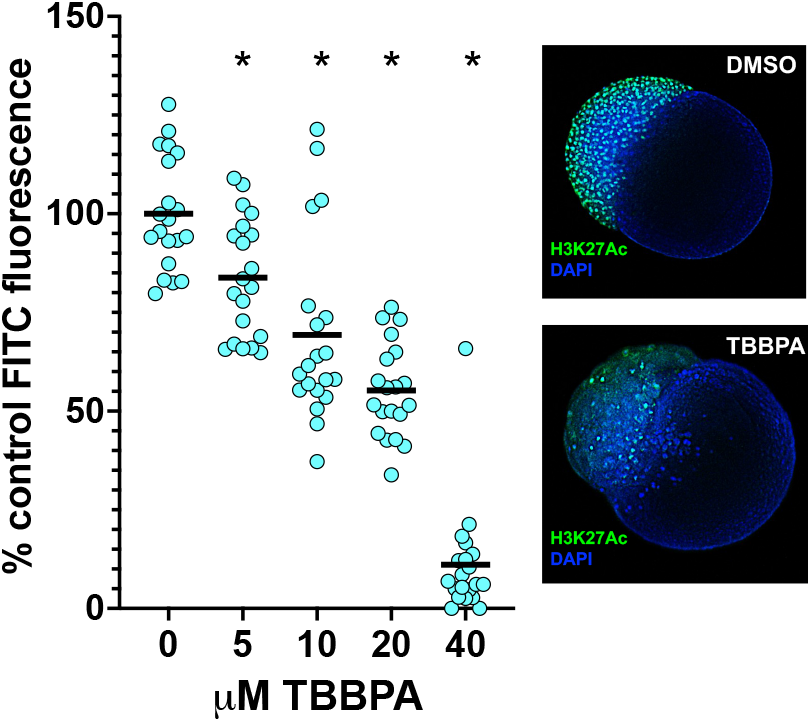
TBBPA reduces H3K27Ac levels during ZGA. Embryos were exposed to TBBPA solutions at 0.75 hpf and immunostained with an anti-H3K27Ac antibody at 3.5 hpf (N=20). Representative images on the right.

**Figure 6.**
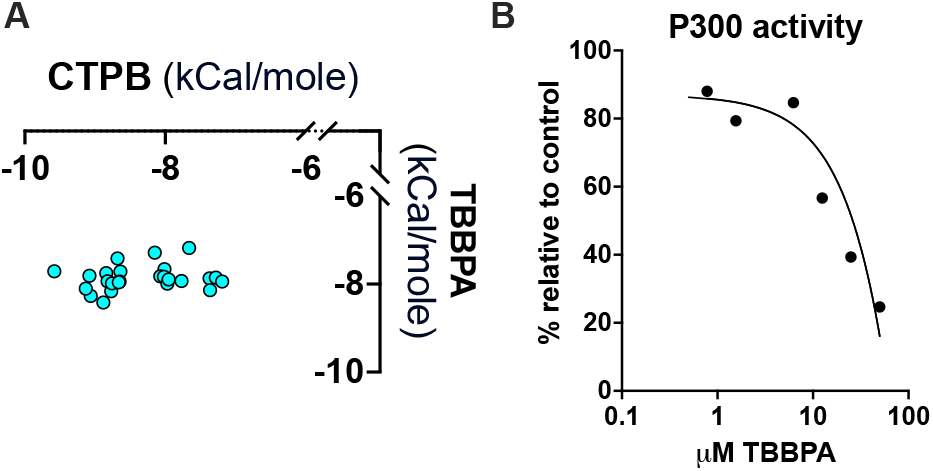
TBBPA potentially targets P300 protein. **(A)** *In silico* molecular docking studies show equivalent binding energy for TBBPA and CTPB (a P300 activator) on the P300 HAT domain. CTPB- (n-(4-chloro-3-trifluoromethyl-phenyl)-2-ethoxy-6-pentadecyl-benzamide is a P300 HAT activator **(B)** TBBPA inhibits P300 activity *in vitro*.

### CTPB co-exposures with TBBPA partially mitigate stage-specific delays of TBBPA exposures alone

Both CTPB-TBBPA co-exposures and pre-exposures partially mitigated TBBPA-induced developmental delays in zebrafish. For co-exposures, while ∼80% of 20 µM TBBPA embryos at ∼3 hpf were at the 128-256 cell stage, only ∼70% of embryos within CTPB co-exposures were at the 512-1k stage, suggesting a mitigation of phenotype **(Fig 7A)**. Partial mitigation of TBBPA-induced developmental delays were also seen with CTPB pre-exposures. While ∼80% of embryos exposed to 20 µM TBBPA exposure were at the sphere cell stage at 6 hpf, 90% of embryos exposed to CTPB prior to TBBPA exposure were at the 30% epiboly stage **(Fig 7B)**. For both co- and pre-exposures, overall CTPB x TBBPA x stage interactions were significant.

**Figure 7.**
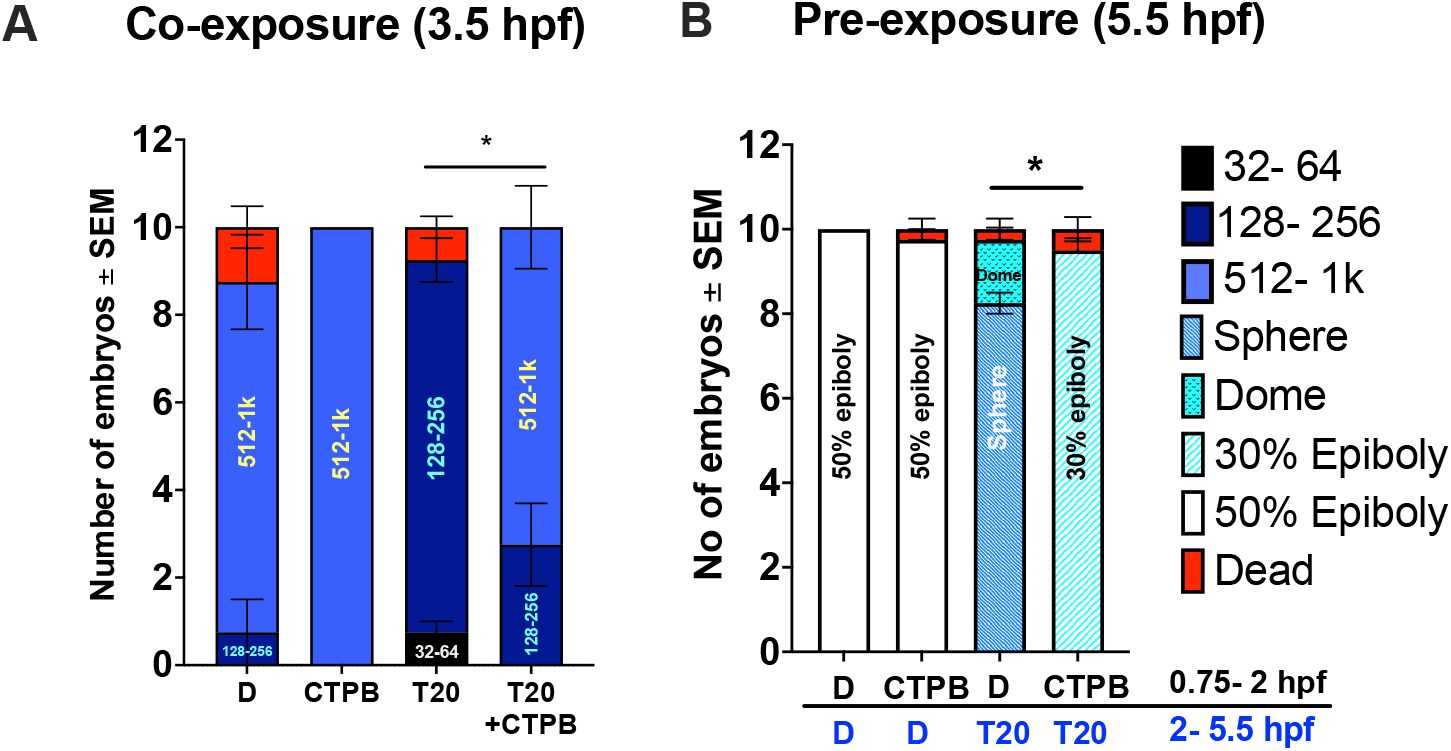
CTPB co- and pre-exposures mitigate TBBPA-induced delays. (A) CTPB-TBBPA co-exposures from 0.75 to 3 hpf partially mitigate TBBPA-induced developmental delays. (B) CTPB pre-exposures from 0.75 to 2 hpf, followed by TBBPA exposures from 2 to 5.5 hpf mitigate TBBPA-induced developmental delays. D= 0.1% DMSO, T20= 20 µM TBBPA, CTPB= 50 µM CTPB. Astrisk (*) denotes statistically significant difference based on 3-way ANOVA followed by Tukey’s test.

## DISCUSSION

TBBPA is a legacy FR that has undergone extensive investigation to identify any adverse health risks. Despite extensive research on TBBPA, understanding its potential effects on embryos during pre-pluripotent stages has been lacking. A prior study showed that aqueous TBBPA exposures led to a substantial embryonic uptake of TBBPA (0.8 ng/g) immediately after exposures were initiated at 2 hpf, suggesting a rapid uptake into cells during pre-pluripotent stages (Liu et al., 2018). Disruptions of these processes can lead to long-term health consequences through persistent genetic and epigenetic insults. Our phenotypic data showed that TBBPA impacted embryonic health during these early stages, with significant developmental delays and mortality during the maternal-to-zygotic transition and zygotic genome activation prior to differentiation. This concurs with previous work in various model organisms demonstrating that blastulation and gastrulation defects can cause embryonic lethality and defects in implantation (Su et al., 2018). Following initial phenotyping, we conducted LPR experiments to assess the long-term impacts of short-term TBBPA exposures. The LPR assay, while a swimming behavioral readout, is an overall indicator of embryonic health. Our LPR data shows that these short-term TBBPA exposures can disrupt LPR at concentrations ranging down to 0.004 µM (4 nM). Within human biomatrices, TBBPA concentrations upto 20 nM in adult plasma (Sunday et al., 2022) and 5.3 nM in cord plasma have been detected, with a study modelling concentration up to 1.2 µM in cord plasma (Sunday et al. 2022). Therefore, LPR hypoactivity was seen at environmentally relevant exposures. Taken together, our phenotypic and behavior data shows that transient TBBPA exposures during pre-pluripotent stages can have short- and long-term impacts on developmental trajectory and function of the embryo.

We then focused on investigating the genetic basis and signature of TBBPA phenotypes during MZT. Since MZT and ZGA are gatekeeping events for further development, insults to these events may be a primary causative factor for the phenotypes observed. We conducted RNA-seq at multiple stages (2, 3.5, and 4.5 hpf) to profile ZGA period and plotted normalized read counts across multiple developmental time points to represent gene expression of maternal, maternal-zygotic and zygotic transcripts. This approach was taken within toxicological studies in previous papers (Dasgupta et al., 2018; Vliet et al., 2018) that identified drivers of epiboly delays. While cumulative read counts obtained from maternal and maternal-zygotic genes showed no differences, zygotic-only (Z) genes showed a significant reduction in expression within TBBPA exposures, demonstrating an inhibition of the ZGA. MZT and ZGA are crucial steps for downstream cell patterning and cell fate determination (Tadros & Lipshitz, 2009), potentially having severe effects on embryo vitality. Overall, these findings support our hypothesis that TBBPA suppresses ZGA-an effect that potentially leads to delay of embryogenesis.

We then further relied on gene ontology assessments to examine the potential initiating mechanism for critical delay. Interestingly, both time-matched and stage-matched comparisons suggested an inhibition of chromatin remodeling, allowing us to focus on histone modifiers as a potential TBBPA target. The histone modification system that is primarily operant during the sensitive time window at which we observe phenotypes is H3K27Ac- (acetylation of Lysine 27 of Histone 3) (Wang et al., 2022). This epigenetic alteration is known to be essential in ZGA, modulating chromatin remodeling and gene expression. Previous work has shown that modification of histone acetylation during embryogenesis can result in neural tube defects and neurodegenerative diseases (Tadros & Lipshitz, 2009). H3K27 acetylation is catalyzed by the P300-CBP protein complex which contains several domains, including a HAT (histone acetyltransferase) catalytic domain that “writes” acetyl groups from acetyl coA onto histones and promotes an open chromatin state that drives gene activation (Dal Piaz et al., 2010). P300 activity is crucial is for early embryonic development, especially during MZT and ZGA (Chan et al., 2019) and studies have shown that a P300 inhibition can result in ZGA failure and developmental arrest (Wang et al., 2022). Based on these, we sought to evaluate whether TBBPA-induced ZGA inhibition and developmental delays were linked to a P300-H3K27Ac inhibitory mechanism. Using immunohistochemistry, we showed that TBBPA exposures indeed inhibit H3K27Ac levels within embryos. Our *in-silico* modeling, combined with *in vitro* P300 assays-both showed that TBBPA potentially targets histone acetyltransferase (HAT) domain of P300. However, whether this was a probably initiating factor *in vivo* remained a crucial question. Therefore, we leveraged CTPB (a small molecular activator of P300), and, through pre- and co-exposure experiments, reliably demonstrated that the presence of CTPB limits TBBPA-induced phenotypes. Collectively, these experiments demonstrated that histone acetylation acts as pivotal mechanism for TBBPA-induced ZGA inhibition, and this process is likely driven by TBBPA acting as a ligand to P300 HAT domain, thereby limiting its HAT activity and H3K27 acetylation, limiting chromatin remodeling and activation of the zygotic genome.

## CONCLUSION

Our data mechanistically shows that TBBPA delays embryonic development and zygotic genome activation by inhibiting P300 and reduces levels of P300-catalyzed histone acetylation. This mode of action is understudied in environmental toxicology, but the results have profound implication for developmental health since histone modifiers (including acetylation) and zygotic genome activation set stage for healthy embryonic development. Our photomotor response data suggests that short term TBBPA exposures can have long term developmental effects even at environmentally relevant concentrations. However, we did not profile molecular responses at these environmentally relevant concentrations since this study was focused on identifying the causative factors of pre-pluripotent development delays through phenotypic anchoring of data. Furthermore, while H3K27Ac is the dominant one, other histone modifiers that are targeted by TBBPA functionally drive phenotypes. Therefore, equipped with data from this work, our future work will-1) profile molecular effects at environmentally relevant concentrations; 2) profile other histone modifiers and 3) conduct chromatin-level studies to understand how TBBPA exposures limit chromatin remodeling and transcription.

## ACKNOWLEDGEMENTS

We thank Mr. John Smink and Clemson Aquatic Animal Research Facility for zebrafish husbandry, Dr. Robyn Tanguay, along with Dr. Lisa Truong and Mr. Ryan Lopez (Oregon State University) for generously providing founder fish for our study, Clemson University Genetics and Bioinformatics Facility for usage of Tapestation, Ms. Maria Baltazar for embryo collection/sorting and Mr. Syed Rubaiyat Ferdous for review of the manuscript.

## SUPPLEMENTAL INFORMATION

Supplemental file contains details for RNA-seq experiments and legends for supplemental tables. Supplemental tables file contains supplemental tables S1 to S8.

## REFERENCES

Akdogan-Ozdilek, B., Duval, K. L. & Goll, M. G. Chromatin dynamics at the maternal to zygotic transition: recent advances from the zebrafish model. F1000Res 9, F1000 Faculty Rev-299 (2020).

Alexander, Jan, Diane Benford, Alan Boobis, and Åke Bergman. “Scientific Opinion on Tetrabromobisphenol A (TBBPA) and its derivatives in food: EFSA Panel on Contaminants in the Food Chain (CONTAM).” EFSA Journal 9, no. 12 (2011): 2477.

(Alexander et al., 2011)Alzualde, A., Behl, M., Sipes, N. S., Hsieh, J.-H., Alday, A., Tice, R. R., Paules, R. S., Muriana, A., & Quevedo, C. (2018). Toxicity profiling of flame retardants in zebrafish embryos using a battery of assays for developmental toxicity, neurotoxicity, cardiotoxicity and hepatotoxicity toward human relevance. Neurotoxicology and Teratology, 70, 40–50. 10.1016/j.ntt.2018.10.002

Cariou, R., Antignac, J.-P., Zalko, D., Berrebi, A., Cravedi, J.-P., Maume, D., Marchand, P., Monteau, F., Riu, A., Andre, F., & bizec, B. L. (2008). Exposure assessment of French women and their newborns to tetrabromobisphenol-A: Occurrence measurements in maternal adipose tissue, serum, breast milk and cord serum. Chemosphere, 73(7), 1036–1041. 10.1016/j.chemosphere.2008.07.084

Chan, S. H., Tang, Y., Miao, L., Darwich-Codore, H., Vejnar, C. E., Beaudoin, J.-D., Musaev, D., Fernandez, J. P., Benitez, M. D. J., Bazzini, A. A., Moreno-Mateos, M. A., & Giraldez, A. J. (2019). Brd4 and P300 Confer Transcriptional Competency during Zygotic Genome Activation. Developmental Cell, 49(6), 867-881.e8. 10.1016/j.devcel.2019.05.037

Cho, J.-H., Lee, S., Jeon, H., Kim, A. H., Lee, W., Lee, Y., Yang, S., Yun, J., Jung, Y.-S., & Lee, J. (2020). Tetrabromobisphenol A-Induced Apoptosis in Neural Stem Cells Through Oxidative Stress and Mitochondrial Dysfunction. Neurotoxicity Research, 38(1), 74–85. 10.1007/s12640-020-00179-z

Dal Piaz, F., Tosco, A., Eletto, D., Piccinelli, A. L., Moltedo, O., Franceschelli, S., Sbardella, G., Remondelli, P., Rastrelli, L., Vesci, L., Pisano, C., & De Tommasi, N. (2010). The identification of a novel natural activator of p300 histone acetyltranferase provides new insights into the modulation mechanism of this enzyme. Chembiochem: A European Journal of Chemical Biology, 11(6), 818–827.

Dasgupta, S., Cheng, V., Vliet, S. M. F., Mitchell, C. A. & Volz, D. C. Tris(1,3-dichloro-2-propyl) Phosphate Exposure During the Early-Blastula Stage Alters the Normal Trajectory of Zebrafish Embryogenesis. Environ Sci Technol 52, 10820–10828 (2018).

Kimmel, C. B.; Ballard, W. W.; Kimmel, S. R.; Ullmann, B.; Schilling, T. F. Stages of embryonic development of the zebrafish. Dev. Dyn. 1995, 203 (3), 253−310.

Mantelingu, K., Kishore, A. H., Balasubramanyam, K., Kumar, G. V. P., Altaf, M., Swamy, S. N., Selvi, R., Das, C., Narayana, C., Rangappa, K. S., & Kundu, T. K. (2007). Activation of p300 Histone Acetyltransferase by Small Molecules Altering Enzyme Structure: Probed by Surface-Enhanced Raman Spectroscopy. The Journal of Physical Chemistry B, 111(17), 4527–4534. 10.1021/jp067655s

Oral, D., Balci, A., Chao, M.-W., & Erkekoglu, P. (2021). Toxic Effects of Tetrabromobisphenol A: Focus on Endocrine Disruption. Journal of Environmental Pathology, Toxicology and Oncology, 40(3). 10.1615/JEnvironPatholToxicolOncol.2021035595

Reed, J. M., Spinelli, P., Falcone, S., He, M., Goeke, C. M., & Susiarjo, M. (n.d.). Evaluating the Effects of BPA and TBBPA Exposure on Pregnancy Loss and Maternal–Fetal Immune Cells in Mice. Environmental Health Perspectives, 130(3), 037010. 10.1289/EHP10640

Shi, Z.-X., Wu, Y.-N., Li, J.-G., Zhao, Y.-F., & Feng, J.-F. (2009). Dietary Exposure Assessment of Chinese Adults and Nursing Infants to Tetrabromobisphenol-A and Hexabromocyclododecanes: Occurrence Measurements in Foods and Human Milk. Environmental Science & Technology, 43(12), 4314–4319. 10.1021/es8035626

Su X, Wu C, Ye X, Zeng M, Zhang Z, Che Y, Zhang Y, Liu L, Lin Y, Yang R. Embryonic lethality in mice lacking Trim59 due to impaired gastrulation development. Cell Death Dis. 2018 Feb 21;9(3):302. doi: 10.1038/s41419-018-0370-y. PMID: 29467473; PMCID: PMC5833458.

Sunday, O. E. et al./person-group>. Review of the environmental occurrence, analytical techniques, degradation and toxicity of TBBPA and its derivatives. Environ Res 206, 112594 (2022).

Tadros, W., & Lipshitz, H. D. (2009). The maternal-to-zygotic transition: A play in two acts. Development, 136(18), 3033–3042. 10.1242/dev.033183

Vliet SM, Dasgupta S, Volz DC. Niclosamide Induces Epiboly Delay During Early Zebrafish Embryogenesis. Toxicol Sci. 2018 Dec 1;166(2):306–317. doi: 10.1093/toxsci/kfy214. PMID: 30165700.

Wang, M., Chen, Z., & Zhang, Y. (2022). CBP/p300 and HDAC activities regulate H3K27 acetylation dynamics and zygotic genome activation in mouse preimplantation embryos. The EMBO Journal, 41(22), e112012. 10.15252/embj.2022112012

Wu, S., Ji, G., Liu, J., Zhang, S., Gong, Y., & Shi, L. (2016). TBBPA induces developmental toxicity, oxidative stress, and apoptosis in embryos and zebrafish larvae (Danio rerio). Environmental Toxicology, 31(10), 1241–1249. 10.1002/tox.22131

Zhang, H., Liu, W., Chen, B., He, J., Chen, F., Shan, X., Du, Q., Li, N., Jia, X., & Tang, J. (2018). Differences in reproductive toxicity of TBBPA and TCBPA exposure in male Rana nigromaculata. Environmental Pollution, 243, 394–403. 10.1016/j.envpol.2018.08.086

Zhou, X., Guo, J., Zhang, W., Zhou, P., Deng, J., & Lin, K. (2014). Tetrabromobisphenol A contamination and emission in printed circuit board production and implications for human exposure. Journal of Hazardous Materials, 273, 27–35. 10.1016/j.jhazmat.2014.03.003

