## Supplemental File for "Flame retardant tetrabromobisphenol A (TBBPA) disrupts histone acetylation during zebrafish maternal-to-zygotic transition"

1. **RNA-seq details**

**Library Construction, Quality Control and Sequencing**

Messenger RNA was purified from total RNA using poly-T oligo-attached magnetic beads. After fragmentation, the first strand cDNA was synthesized using random hexamer primers, followed by the second strand cDNA synthesis using either dUTP for directional library or dTTP for non-directional library[1].

For the non-directional library, it was ready after end repair, A-tailing, adapter ligation, size selection, amplification, and purification. The library was checked with Qubit and real-time PCR for quantification and bioanalyzer for size distribution detection. Quantified libraries will be pooled and sequenced on Illumina platforms, according to effective library concentration and data amount.

**Bioinformatics**

***Data Quality Control***

Raw data (raw reads) of fastq format were firstly processed through in-house perl scripts. In this step, clean data (clean reads) were obtained by removing reads containing adapter, reads containing ploy-N and low quality reads from raw data. At the same time, Q20, Q30 and GC content the clean data were calculated. All the downstream analyses were based on the clean data with high quality.

***Reads mapping to the reference genome***

Reference genome and gene model annotation files were downloaded from genome website directly. Index of the reference genome was built using Hisat2 v2.0.5 and paired-end clean 1 reads were aligned to the reference genome using Hisat2 v2.0.5. We selected Hisat2[2] as the mapping tool for that Hisat2 can generate a database of splice junctions based on the gene model annotation file and thus a better mapping result than other non-splice mapping tools.

***Quantification of gene expression level***

featureCounts^[3]^ v1.5.0-p3 was used to count the reads numbers mapped to each gene. And then FPKM of each gene was calculated based on the length of the gene and reads count mapped to this gene. FPKM, expected number of Fragments Per Kilobase of transcript sequence per Millions base pairs sequenced, considers the effect of sequencing depth and gene length for the reads count at the same time, and is currently the most commonly used method for estimating gene expression levels.

***Differential expression analysis***

Differential expression[4,5] analysis of two conditions/groups (two biological replicates per condition) was performed using the DESeq2Rpackage (1.20.0). DESeq2 provide statistical routines for determining differential expression in digital gene expression data using a model based on the negative binomial distribution. The resulting P-values were adjusted using the Benjamini and Hochberg's approach for controlling the false discovery rate . Genes with an adjusted P-value

<=0.05found by DESeq2 were assigned as differentially expressed. (For edgeR[6] without biological replicates) Prior to differential gene expression analysis, for each sequenced library, the read counts were adjusted by edgeR program package through one scaling normalized factor. Differential expression analysis of two conditions was performed using the edgeR R package (3.22.5). The P values were adjusted using the Benjamini & Hochberg method. Corrected P-value of 0.05 and absolute foldchange of 2were set as the threshold for significantly differential expression.

2. **Supplemental Table legends**

**Tables S1 and S2:** Raw data for LPR studies from 2 plates- plates 1 and 2. Yellow highlight- embryo number; blue highlight- time bin number. Blue text with * indicates embryo was excluded due to mortality or abnormality.

**Tables S3 and S4:** FPKM read counts and differential expressions following RNA-seq at 2 hpf (Table S3) and 3.5 and 4.5 hpf (Table S4).

**Tables S5.** Gene Ontology for all differentially expressed genes for time-matched comparison between TBBPA-3.5 hpf and DMSO-3.5 hpf samples. DEG filter: |log2FC|>1, p*_adj_*<0.05

**Tables S6.** Gene Ontology for differentially expressed genes with negative fold changes for time-matched comparison between TBBPA-3.5 hpf and DMSO-3.5 hpf samples. DEG filter: log2FC<-1, p*_adj_*<0.05

**Tables S7.** Gene Ontology for all differentially expressed genes for stage-matched comparison between TBBPA-4.5 hpf and DMSO-3.5 hpf samples. DEG filter: |log2FC|>1, p*_adj_*<0.05

**Tables S8.** Gene Ontology for all differentially expressed genes with negative fold changes for stage-matched comparison between TBBPA-4.5 hpf and DMSO-3.5 hpf samples. DEG filter: log2FC<-1, p*_adj_*<0.05.

**Table S9.** List of maternal, zygotic and maternal-zygotic genes obtained from Bhat et al 2023 paper.

**Table S10.** List of available crystals structures of p300 (UniProtKB: Q09472) as on 2023-06-04. Quality of models available in PDB and PDB-REDO repositories are shown. Model geometry and fit model/data are scored on an arbitrary scale of -2 (worst) to 2 (best) based on the wwPDB validation report. Bound ligand information is also given.

**Table S11.** Estimated binding energies of the compounds with the catalytic domain of p300.

**Table S12.** Estimated binding energies of the compounds with the bromodomain of p300.

3. **References**
1. Parkhomchuk D, Borodina T, Amstislavskiy V, et al. Transcriptome analysis by strand-specific sequencing of complementary DNA[J]. Nucleic acids research, 2009, 37(18): e123-e123.
2. Mortazavi A, Williams B A, McCue K, et al. Mapping and quantifying mammalian transcriptomes by RNA-Seq[J]. Nature methods, 2008, 5(7): 621-628.
3. Liao Y1, Smyth GK, Shi W. featureCounts: an efficient general purpose program for assigning sequence reads to genomicfeatures.Bioinformatics.2014 ,30(7):923-30.
4. Love M I, Huber W, Anders S. Moderated estimation of fold change and dispersion for RNA-seq data with DESeq2[J]. Genome biology, 2014, 15(12): 1-21.
5. Anders S, Huber W. Differential expression analysis for sequence count data[J]. Genome biol, 2010, 11(10): R106.
6. Robinson M D, McCarthy D J, Smyth G K. edgeR: a Bioconductor package for differential expression analysis of digital gene expression data[J]. Bioinformatics, 2010, 26(1): 139-140.
